# A network of MAP-Kinase pathways and transcription factors regulates cell-to-cell communication and cell wall integrity in *Neurospora crassa*

**DOI:** 10.1101/281931

**Authors:** Monika S. Fischer, Vincent W. Wu, Ji E. Lee, Ronan C. O’Malley, N. Louise Glass

**Affiliations:** The Plant and Microbial Biology Department, The University of California, Berkeley, CA 94720-3102; The Energy Biosciences Institute, The University of California, Berkeley, CA 94720-3102; Environmental Genomics and Systems Biology Division, Lawrence Berkeley National Laboratory, Berkeley, CA 94720; U.S. Department of Energy, Joint Genome Institute, Walnut Creek, CA 94598, USA

**Author notes:** **Corresponding Author** N. Louise Glass University of California, Berkeley Department of Plant & Microbial Biology 341A Koshland Hall Berkeley, California 94720.

**Keywords:** *Neurospora crassa*, MAP Kinase, Regulatory Networks, DAP-seq, cell-to-cell communication, cell fusion

## Abstract

Maintenance of cell integrity and cell-to-cell communication are fundamental biological processes. Filamentous fungi, such as *Neurospora crassa*, depend on communication to locate compatible cells, coordinate cell fusion, and establish a robust hyphal network. Two MAP-Kinase pathways are essential for communication and cell fusion in *N. crassa*; the Cell Wall Integrity/MAK-1 pathway and the MAK-2 (signal response) pathway. Previous studies have demonstrated several points of cross talk between the MAK-1 and MAK-2 pathways, which is likely necessary for coordinating chemotropic growth toward an extracellular signal, and then mediating cell fusion. Canonical MAP-Kinase pathways begin with signal reception and end with a transcriptional response. Two transcription factors, ADV-1 and PP-1, are essential for communication and cell fusion. PP-1 is the conserved target of MAK-2, while it is unclear what targets ADV-1. We did RNAseq on *Δadv-1, Δpp-1*, and wild-type cells and found that ADV-1 and PP-1 have a shared regulon including many genes required for communication, cell fusion, growth, development, and stress response. We identified ADV-1 and PP-1 binding sites across the genome by adapting the *in vitro* method of DNA-Affinity Purification sequencing (DAP-seq) for *N. crassa*. To elucidate the regulatory network, we misexpressed each transcription factor in each upstream MAPK deletion mutant. Misexpression of *adv-1* was sufficient to fully suppress the phenotype of the *Δpp-1* mutant and partially suppress the phenotype of the *Δmak-1* mutant. Collectively, our data demonstrate that the MAK-1-ADV-1 and MAK-2- PP-1 pathways form a tight regulatory network that maintains cell integrity and mediates communication and cell fusion.

## INTRODUCTION

Cell-to-cell communication is a fundamental biological process across the tree of life. There is abundant work detailing mechanisms that mediate communication processes in diverse organisms, such as quorum sensing in bacteria, neurotransmission in mammals, or pheromone sensing in *Saccharomyces cerevisiae* (Perbal 2003; Merlini *et al.* 2013). All of these systems share a general mechanism of signal release and reception that subsequently initiates specific molecular responses that often lead to changes in transcription.

Filamentous fungi, such as *Neurospora crassa*, depend on cell-to-cell communication to locate compatible cells and coordinate the process of cell fusion. *N. crassa* is a well-developed model for investigating mechanisms of cell-cell communication, chemotropic growth, and cell fusion at several points throughout the life cycle. During germination of genetically identical asexual spores, individual cells (germlings) collaborate to establish a new colony by engaging in cell-to-cell communication and chemotropic growth that ultimately results in cell fusion (Glass 2004; Fleissner *et al.* 2009; Richard *et al.* 2012; Leeder *et al.* 2013; Bastiaans *et al.* 2015). Germling fusion is an important aspect of colony establishment, and as a mature colony develops, hyphae within a colony also undergo chemotropic growth and fusion to further reinforce a robust hyphal network (Hickey *et al.* 2002). Hyphal fusion also occurs between different colonies, which is important for mediating post-fusion self/non-self recognition mechanisms and heterokaryosis (Garnjobst and Wilson 1952; Saupe 2000; Glass and Kaneko 2003; Simonin *et al.* 2012). Cell-to-cell communication and fusion are also necessary for fertilization during the process of sexual reproduction. In filamentous ascomycete species, such as *N. crassa,* fertile hyphae (trichogynes) from protoperithecia chemotropically grow towards the male gamete with the goal of cell fusion and the initiation of perithecium development, karyogamy, meiosis, and finally ascospore development (Kim and Borkovich 2004; Engh *et al.* 2010).

More than 60 genes have been identified as required for the process of germling communication and cell fusion (Fu *et al.* 2011; Read *et al.* 2012; Palma-Guerrero *et al.* 2013; Dettmann *et al.* 2014; Fleißner and Herzog 2016). Stains carrying mutations in many of the genes involved in mediating germling or hyphal fusion show a pleiotropic vegetative and female sexual development phenotype. For example, in both *N. crassa* and *Sordaria macrospora*, most fusion mutants fail to develop female reproductive structures (protoperithecia) (Engh *et al.* 2010; Fu *et al.* 2011). Some of these genes are also essential for sexual reproduction, but many are not, indicating that asexual (germling and hyphal) cell fusion is mechanistically distinct from sexual (gamete) fusion. Among the >60 known genes, there are many encoding hypothetical proteins of unknown function, but also genes/proteins in characterized pathways, including two ERK-like Mitogen Activated Protein Kinase (MAPK) pathways and pathways involved in the production of reactive oxygen species (ROS), actin organization, and calcium signaling.

Two of the three predicted MAPK pathways in *N. crassa* are required for germling communication and cell fusion (Colot *et al.* 2006; Park *et al.* 2008). In general, MAPK pathways are intermediaries that transduce information from one part of the cell (i.e. a cell-surface sensor; input) to factors in the nucleus that alter gene transcription (output). The two MAPK pathways essential for communication and cell fusion are the MAK-2 pathway, which is homologous to the pheromone response pathway in *S. cerevisiae*, and the MAK-1 pathway, which is homologous to the cell wall integrity pathway in *S. cerevisiae*. In *N. crassa,* the core conserved components of the MAK-1 pathway are MIK-1 (MAPKKK), MEK-1 (MAPKK), MAK-1 (MAPK), and SO, a scaffold important for some functions of the MAK-1 pathway (Teichert *et al.* 2014; Weichert *et al.* 2016). It is currently unknown which transcription factors are targets of the MAK-1 pathway, although transcription of several genes is dependent on MAK-1, and the rhythmic pattern of expression for a few MAK-1-dependent genes mirrors the rhythmic expression pattern of ADV-1-dependent genes (Bennett *et al.* 2013; Dekhang *et al.* 2017). Additionally, both *mak-1* and *adv-1* are targets of the circadian clock in *N. crassa* (Bennett *et al.* 2013; Dekhang *et al.* 2017). The core components of the conserved *N. crassa* MAK-2 pathway are NRC-1 (MAPKKK), MEK-2 (MAPKK), MAK-2 (MAPK), and HAM-5, a scaffold protein important during cell fusion (Dettmann *et al.* 2014; Jonkers *et al.* 2014). PP-1 is the predicted downstream transcription factor target of MAK-2; a microarray expression study demonstrated that there is overlap between MAK-2-dependent and PP-1-dependent gene expression (Leeder *et al.* 2013). Several other studies have documented cross talk between the MAK-1 and MAK-2 pathways in *N. crassa* and in other filamentous ascomycete fungi, some of which is mediated by the STRIPAK complex (Maerz *et al.* 2008; Maddi *et al.* 2012; Dettmann *et al.* 2012, 2013; Leeder *et al.* 2013; Fu *et al.* 2014).

Transcription is a common point of regulation for biological processes, and two conserved transcription factors in *N. crassa* (ADV-1 and PP-1) are essential for germling communication and fusion. Both the *Δadv-1* and *Δpp-1* deletion mutants have a pleiotropic phenotype and were initially identified as female-sterile mutants (Li 2005; Colot *et al.* 2006; Fu *et al.* 2011). PP-1 is a C2H2 zinc finger transcription factor that is homologous to the *S. cerevisiae* pheromone response pathway transcription factor, *STE12* (Leeder *et al.* 2013). The core component of this pathway in *S. cerevisiae* is the Fus3p MAPK cascade, that once activated, leads to de-repression of Ste12p (Merlini *et al.* 2013). Ste12-like proteins, including PP-1, have two C2H2-Zn^2+^motifs and a homeobox-like STE domain involved in binding DNA (Errede and Ammerer 1989). In *N. crassa* the STE domain is essential for PP-1 function, but the C2H2-Zn^2+^motifs are dispensable (Leeder *et al.* 2013). Ste12-like transcription factors in fungi are regulated by direct phosphorylation and phosphorylation of associated regulatory proteins (Blackwell *et al.* 2007). Several phosphorylation sites have been identified on PP-1 in *N. crassa*, but the biological significance of these sites remains unknown (Leeder *et al.* 2013; Xiong *et al.* 2014; Jonkers *et al.* 2014).

ADV-1 is a Zn(II)_2_Cys_6_ transcription factor that, like PP-1, regulates growth, and asexual and sexual development. ADV-1 is not as broadly conserved as PP-1, and clear *adv-1* homologs are restricted to the filamentous Ascomycete species (Pezizomycotina). In the self-fertile (homothallic) species *S. macrospora*, the *adv-1*-ortholog, *pro1*, is required for protoperithecial development, while heterothallic *N. crassa* does not require *adv-1* for protoperithecia development (Masloff *et al.* 1999). In *N. crassa, adv-1* is essential for post-mating perithecial development, asexual cell fusion, and wild-type-like growth rate. Both ADV-1 and Pro1 have a GAL4-like DNA-binding domain and a transcription-activation domain. In contrast to the *S. cerevisiae* protein GAL4p, Pro1 lacks a coiled-coil dimerization domain, indicating that Pro1 (and ADV-1) likely function independently (Masloff *et al.* 2002). Unlike PP-1, upstream factors that influence ADV-1 regulated transcription are largely unknown, with the exception of a ChIPseq experiment that identified *adv-1* as a target of the circadian clock master regulator, the White Collar Complex (Smith *et al.* 2010). ADV-1 is essential for developmental oscillations (i.e. conidiation) and the quantity of *adv-1* mRNA and ADV-1 protein oscillates with the clock (Smith *et al.* 2010). Furthermore, the expression of ADV-1 target genes in a mature colony matches the rhythm of other clock-controlled genes. These data led to the hypothesis that hyphal fusion is a clock-regulated developmental process (Dekhang *et al.* 2017).

The *Δadv-1* and *Δpp-1* mutants share many phenotypes across filamentous fungi, yet the relationship between these two transcription factors remains unclear. Independent expression profiling studies investigating ADV-1 or PP-1-dependent transcription indicate that at least some genes require ADV-1 and PP-1 for wild type levels of expression (Li 2005; Nowrousian *et al.* 2007; Leeder *et al.* 2013; Dekhang *et al.* 2017). Here, we compared expression profiles using RNAseq on *Δadv-1* and *Δpp-1* germlings relative to expression patterns in wild-type germlings. To identify genes that were regulated and bound by ADV-1 or PP-1, we developed an *in vitro* method for identifying transcription factor binding sites in *N. crassa* and other fungi called DNA Affinity Purification sequencing (DAP-seq). DAP-seq is similar to ChIPseq, except that *in vitro* synthesized transcription factors are incubated with native genomic DNA. DNA fragments bound by the transcription factor are then identified by high throughput sequencing methods. DAP-seq has been used in a global analysis of transcription factor binding sites in *Arabidopsis thaliana* (O’Malley *et al.* 2016). Our data showed that PP-1 directly regulates *adv-*1; ADV-1 is the primary regulator of many genes that are important for asexual growth, cell fusion and development. To investigate the linkage between *mak-1, mak-2, adv-1*, and *pp-1*, we used misexpression and phenotypic analyses. Our data showed that *mak-1* primarily functions upstream of *adv-1*, and *mak-2* primarily functions upstream of *pp-1*. However, the MAK-1/ADV-1 pathway and the MAK-2/PP-1 pathway engage in crosstalk and both pathways form a regulatory network that mediates growth, communication, fusion, and the response to cell wall stress.

## MATERIALS & METHODS

### RNA isolation

Strains used: FGSC 2489 (wild-type), FGSC 11042 (*Δadv-1*) and *Δpp-1* (Leeder *et al.* 2013). Each strain was initially grown on Vogel’s Minimal Medium (VMM) (Vogel 1956) agar slant tubes for 5 days. Conidia harvested, filtered through cheesecloth, and then inoculated into 100mL of liquid VMM in a 250mL flask to a final concentration of 10^6^conidia/mL. Flasks were incubated at 30°C in constant light for 2.5 hours shaking at 220 rpm to induce germination, followed by 2.5 hours stationary incubation to allow communication to occur. *Δpp-1* conidia have slightly delayed germination, thus these conidia were shaken for 3 hours, followed by 2.5 hours stationary incubation. Germlings were harvested by vacuum filtration over nitrocellulose paper and transferred to a 2mL screw cap tube that was immediately frozen with liquid N_2_. Experimental design for each strain: conidia from one VMM slant tube was used to inoculate 8 flasks of liquid VMM, then two flasks were pooled during germling harvest, resulting in a total of 4 samples per strain. RNA was extracted using a previously described TRIzol-based method (Leeder *et al.* 2013). RNA quality and concentration were quantified via Bioanalyzer at the qb3 facility at UC Berkeley. Three samples per strain with the highest quality and concentration of RNA were submitted for library prep and sequencing on an Illumina HiSeq3000 at the UC Davis DNA Technologies Core.

### RNAseq data analysis and visualization

Fast-X Toolkit (http://hannonlab.cshl.edu/fastx_toolkit/index.html) was used to filter out low quality raw reads (∼11-12% of all reads) and Tophat (Langmead *et al.* 2009) mapped high quality reads to the *N. crassa* transcriptome version 12 (ftp://ftp.broadinstitute.org/pub/annotation/fungi/neurospora_crassa/assembly/NC12) Differential expression was calculated using three independent methods: Cuffdiff (Roberts *et al.* 2010), DESeq2 (Love *et al.* 2014), and EdgeR (Robinson *et al.* 2010; McCarthy *et al.* 2012). We defined the threshold for significant differential expression to be −2 < log_2_FoldChange < 2 and p.adj<0.01. RNAseq raw data (.fastq) is available at the NCBI Sequence Read Archive with accession number SRP133239.

We used the Circos data visualization tool (Krzywinski *et al.* 2009) to generate Figure 2. Genes are grouped according to known function or predicted function based on homology (BLASTp, FungiDB, or the Broad Institute’s Fungal Orthogroups Repository v1.1). Highlighted gene ID’s indicate genes that have an ADV-1 binding site or a PP-1 binding site 2kb upstream from the start codon. PP-1 binding sites were identified solely via DAP-seq, whereas ADV-1 binding sites are the consensus between DAP-seq data and a previously published ChIPseq dataset (Dekhang *et al.* 2017).

### Genomic DNA Library Preparation

DAP-seq was originally developed for *A. thaliana* (O’Malley *et al.* 2016). Here we adapted the protocol in O’Malley *et al* for use with *N. crassa*. FGSC 2489 was grown in liquid VMM for 24 hours at 25°C and shaking at 220rpm. Mycelia were harvested using vacuum filtration over Whatman #1 filter papers, and transferred into 2ml tubes for flash freezing in liquid N_2_. Cells were ruptured by bead beating for 1 minute with 1mm silica beads and DNA lysis buffer (0.05M NaOH, 1mM EDTA, 1% TritonX) was added to each sample tube. DNA was purified using DNeasy Blood & Tissue kit (Qiagen Inc.), and sheared to 300bp peak using Covaris LE220 sonicator. AMPure XP beads were used to remove DNA above and below target molecular weight. Initially, sheared DNA was mixed in with AMPure XP beads (in PEG-8000) at a ratio of 100:60. At this ratio, beads bind DNA with molecular weight above 700bp. Supernatant from this primary binding was taken and added to new beads where final ratio of DNA solution to PEG-8000 was at 100:90. At this ratio, DNA below ∼300bp do not bind to AMPure XP beads, and remaining DNA was be eluted for library preparation. KAPA library kit for illumina sequencing was used to prepare final libraries and stored at −20°C for later use.

### Transcription factor cloning, transcription, translation and DNA Affinity Purification (DAP)

TF open reading frames (ORF) were amplified from cDNA using RNA to cDNA ecodry premix (Clonetech). Amplified TFs sequences were inserted into pIX *in vitro* expression vector modified to contain an N-termainl HALO-Tag (O’Malley *et al.* 2016). Vector backbone was amplified and assembled with TF ORFs using Gibson assembly and then transformed into competent *E. coli* cells for storage and production.

In vitro transcription and translation of TFs was achieved by using Promega TnT T7 Rabbit Reticulocyte Quick Coupled Transcription/Translation System. 1µg of plasmid DNA, 60µl of TnT Master Mix, and 1.5µl of 1mM methionine were combined and incubated overnight at room temperature. Expression was verified using Western blot analysis with Promega HaloTag monoclonal antibody. Completed TnT reactions were incubated with 20ng of genomic DNA library, 1µg salmon sperm for blocking, and 20µl Promega Magne HaloTag Beads on a rotator for 1 hour at room temperature. Bead bound proteins and protein bound DNA were washed three times with 2.5% Tween20 in PBS. HaloTag beads were resuspended in 30µl ddH_2_O and heated to 98°C for 10 minutes to denature protein and release DNA fragments into solution. Supernatant was transferred to a new tube for PCR amplification. DNA was amplified for final libraries using KAPA Hifi polymerase for 12-16 cycles of PCR.

### DAP-seq data analysis

Filtered reads were mapped against *N. crassa* OR74A genome (v12) using bowtie2 v2.3.2 (Kim *et al.* 2013). SamTools (Li *et al.* 2009) was used to convert .sam to .bam files and to create .bai index files for viewing reads on IGV (Integrated Genomics Viewer). MACS2 (Zhang *et al.* 2008) with p-value cutoff at 0.001 was used for calling peaks. A negative control data set consisting of DAP pull-down with Promega TnT master mix with no plasmid added, salmon sperm, and genomic DNA was also input into MACS2 as the control condition. We repeated the ADV-1 DAP-seq once and pooled the results of both DAP-seq runs. DAP-seq data is available at the NCBI Sequence Read Archive with accession number SRP133627.

### Motif construction

To construct biologically meaningful transcription factor DNA binding motifs, we used DAP-seq peaks associated with differentially expressed genes according to corresponding RNAseq data. Sequences of these true positive peaks were collected using a custom python script that reads in genomic position of each peak from the MACS2 output file, and ascertains the sequence from *N. crassa* OR74A Broad v12 genome FASTA file. The script output is a FASTA file with sequences from all true positive peaks. Output FASTA files were input into MEME or DREME v4.12.0 with flags maxw =20, minsites = 5, nmotifs = 8, denoting max width of motif, minimum number of sites for each motif and number of motifs to generate respectively. For ADV-1 specifically, we used set of 41 DAP binding peak sequences. These peaks were within the promoter regions of from genes that fit two parameters: within 1kbp upstream of the ATG start site according to DAP-seq, as well as 4 fold down-regulated in the *Δadv-1* mutant as compared to wild-type. Nucleotide sequences for these 41 peaks were fed into MEME v4.12.0 to build the ADV-1 binding motif. For PP-1, we loosened the parameters slightly to 2 fold down-regulated in *Δpp-1* as compared to wild-type, due to the small number of genes both directly bound according to DAP-seq and differentially expressed at a 4 fold level. Nucleotide sequences for 22 peaks were fed into MEME v4.12.0 to build the PP-1 binding motif.

### qRT-PCR

Germlings were prepared and RNA extracted as described for RNAseq samples. qRT-PCR reactions were prepared following the manufacturer protocols for the Bioline SensiFast(tm) SYBR^®^ no-ROX One-Step kit and Bio-Rad CFX Connect™ Real-Time PCR Detection System. Each sample was replicated at least 4 times within a 96-well-plate and total reaction volume was 20µL. Expression data was normalized to both actin and wild-type following the 2^−ΔΔCt^ method (Livak and Schmittgen 2001).

### Strain construction

Misexpression strains were made by transforming the *his-3*-targeted vectors described below into *his-3*^*-*^ deletion strains as previously described (Colot *et al.* 2006). Positive transformants were selected for histidine prototrophy and hygromycin resistance. To avoid any off-target affects that may have resulted from the transformation process, we backcrossed each transformant to *his-3* (FGSC 9716 or FGSC 6103). Transformants that were incapable of going through a cross were purified via microconidial isolation (Pandit and Maheshwari 1994). All genotypes were confirmed via PCR.

The *Ptef1-adv-1-v5 his-3* vector was constructed by amplifying *adv-1* from genomic DNA with primers that omitted the native STOP codon and added *Xba*I and *Pac*I cut sites on either end of the *adv-1* ORF. PCR products were gel purified and blunt ligated into pCR^®^-Blunt. The *adv-1* sequence was ligated into an in-house vector containing V5 using *Xba*I and *Pac*I sites. *adv-1-v5* was amplified using a reverse primer that added a TGA stop codon at the end of the V5 sequence, in addition to *Eco*RI and *Apa*I cut sites. This PCR product was gel purified and blunt ligated into pCR^®^-Blunt, digested and ligated into an in-house vector based on pMF272 (Freitag *et al.* 2004) containing the *tef-1* promoter (*Ptef1*) using *Xba*I and *Apa*I cut sites. The *Ptef1-pp-1-v5* vector was constructed by amplifying *pp-1* from the *Pccg1-pp-1-gfp his-3* vector (Leeder *et al.* 2013) with primers that added *Bam*HI and *Pac*I sites. This PCR product was ligated into pCR^®^-Blunt, digested and ligated into the *Ptef1-adv1-V5 his3* vector using *Bam*HI and *Pac*I cut sites, which resulted in replacement of the *adv-1* sequence with *pp-1* coding sequences. The sequence of *Ptef1-adv1-v5* and *Ptef1-pp-1-v5* in the *his-3* vectors was confirmed via Sanger sequencing prior to transforming into *his3*^*-*^ deletion strains.

### Growth, fusion, mating, and cell wall stress assays

Standard *N. crassa* growth conditions, media, and protocols are available at http://www.fgsc.net/Neurospora/NeurosporaProtocolGuide.htm. For assays to quantify aerial hyphae height, growth rate, and germling communication, conidia were grown up and harvested as described for RNAseq. Conidial suspensions were diluted to specified concentrations and immediately used for phenotyping assays. Aerial hyphae were grown up from 10^6^ conidia in 1mL of liquid VMM, incubated at 30°C in the dark for 72 hrs (n=6). Growth rate was measured by inoculating 100µL of 10^6^conidia/mL onto one end of a glass race tube containing 25mL of VMM 1.5% agar. Linear growth in race tubes was measured every 24 hrs for at least 4 days (n=3). Germling communication assays consisted of spreading 300µL of 10^7^conidia/mL on a 9 cm VMM 1.5% agar plate. Plates were incubated at 30°C in the dark for 3.5hours. To visualize germlings, agar squares (∼1cm^2^) were excised and observed with a Zeiss Axioskop 2 equipped with a Q Imaging Retiga-2000R camera (Surrey) using a 40x/1.30 Plan-Neofluar oil immersion objective and the iVision Mac4.5 software (Heller *et al.* 2016). Germling communication frequency was determined with the ImageJ Cell Counter tool (http://imagej.nih.gov/ij/). For mating assays, female strains were inoculated onto Synthetic Cross (SC) agar 5.4cm plates (Westergaard and Mitchell 1947). These plates were incubated at 30°C in the dark for two days, and then moved to 25°C and light for an additional 5 days to allow for full production of protoperithecia. Male strains were grown up on VMM as described above. Mating was initiated by inoculating female plates with 150µL of un-diluted conidial suspension of male strains of the opposite mating type. Cell wall stress assays were conducted on large petri plates containing 45mL of VMM with 1.5% agar and 2% FGS. The following cell wall stress drugs were added and mixed to VMM+FGS immediately prior to pouring each plate (one drug/plate): 1.3µg/mL Caspofungin, 1.5mg/mL Calcofluor White, and 1mg/mL Congo Red. Conidia for cell wall stress tests were grown up and harvested as described above, then diluted to 10^6^spores/mL. A 1:5 dilution series was prepared starting with 10^6^spore/mL as the most concentrated dilution. Conidial solutions were then spotted onto freshly poured plates at 5µL/spot.

## RESULTS

### ADV-1 and PP-1 have a shared regulon in germlings

To investigate the how ADV-1 (<u>NCU07392</u>) and PP-1 (<u>NCU00340</u>) regulate germling communication and fusion, we compared expression profiles of *Δadv-1, Δpp-1*, and wild-type germlings. We extracted RNA from germlings 2.5 hours after germination, which is when the majority of wild-type germlings were actively engaging in chemotropic growth and cell fusion (Figure 1A). Our RNAseq data confirm previous studies that implicated PP-1 and ADV-1 as transcriptional activators (Masloff *et al.* 2002; Leeder *et al.* 2013). Figure 1B illustrates that the vast majority of differentially expressed genes were down regulated in *Δadv-1* and *Δpp-1* mutants compared to wild-type germlings (Figure 1B). We defined significant differential expression in each mutant compared to wild-type as - 2>log_2_FoldChange>2, adjusted p-value<0.01, and consensus among three different programs that calculate differential expression; DESeq2, EdgeR, and Cuffdiff (Files S1 and S2).

**Figure 1.**
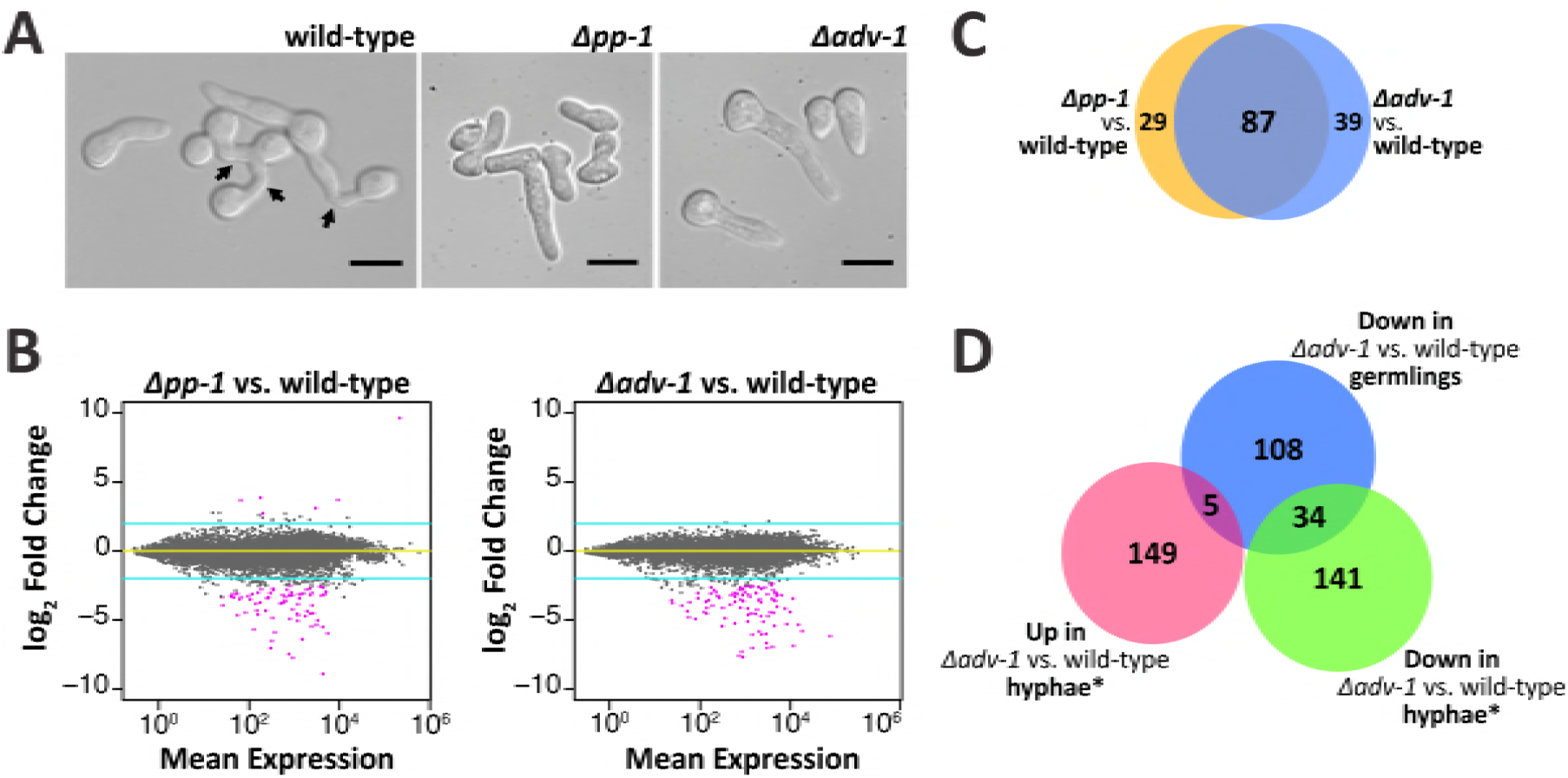
ADV-1 and PP-1 have a shared regulon in germlings. **(A)** Microscopic images showing germling morphology at the time point when we extracted RNA for RNAseq. Arrows indicate fusion events. **(B)** MA plots depicting total RNAseq data for each transcription factor mutant versus the parental wild-type strain. Turquoise lines denote threshold of 2<log_2_FC<-2, and pink points indicate genes with significant differential expression (p<0.01, DESeq2). **(C)** Number of genes significantly down regulated in each mutant as compared to the wild-type parental strain (consensus between CuffDiff, EdgeR, and DESeq2, log_2_FC<-2 and adjusted p-value<0.01). Blue circle is *Δadv-1* compared to wild-type germlings, orange circle is *Δpp-1* compared to wild-type germlings. **(D)** Number of significantly differentially expressed genes in *Δadv-1 as* compared to the parental wild-type strain in hyphae* compared to germlings (− 2>log_2_FC>2 and p<0.01, Cuffdiff). *Hyphal data from Dekhang *et al.* 2017 where RNAseq data was collected from three different time points. Genes were included if they were differentially expressed during at least one time point.

We did not identify any significantly up regulated genes in the *Δadv-1* mutant as compared to wild-type germlings; 17 genes were up regulated in *Δpp-1* cells as compared to wild-type germlings (File S1). The *a*-pheromone precursor gene, *mfa-1*, was the most highly expressed gene in *Δpp-1* (*mat a*) germlings (File S1); it is a clear outlier in the top right corner of Figure 1B. These data complement a previous study that used qRT-PCR to show that both *ccg-4* (*A*-pheromone precursor) and *mfa-1* are substantially over-expressed in *Δpp-1* cells as compared to wild-type (Leeder *et al.* 2013), indicating that PP-1 specifically represses expression of the mating pheromones in germlings.

Analyses of RNAseq data identified 155 significantly down regulated genes in either *Δadv-1* or *Δpp-1* as compared to wild-type germlings (Figure 2 and File S2). There was substantial overlap between the down regulated genes in both mutants (Figure 1C). Of the down regulated genes in *Δpp-1* germlings, 75% (87/116) were also down regulated in *Δadv-1* cells, while 69% (87/126) of the genes down regulated in *Δadv-1* germlings were also down regulated in *Δpp-1* cells. Before imposing a threshold of significant differential expression, we first calculated the distance between samples on our entire RNAseq dataset. These data further demonstrated that gene expression was more similar between *Δadv-1* and *Δpp-1* germlings than either mutant compared to wild-type germlings (Figure S1). We also did not observe any pattern in the genomic location of genes that are regulated by ADV-1 and/or PP-1 (Figure S2).

**Figure 2.**
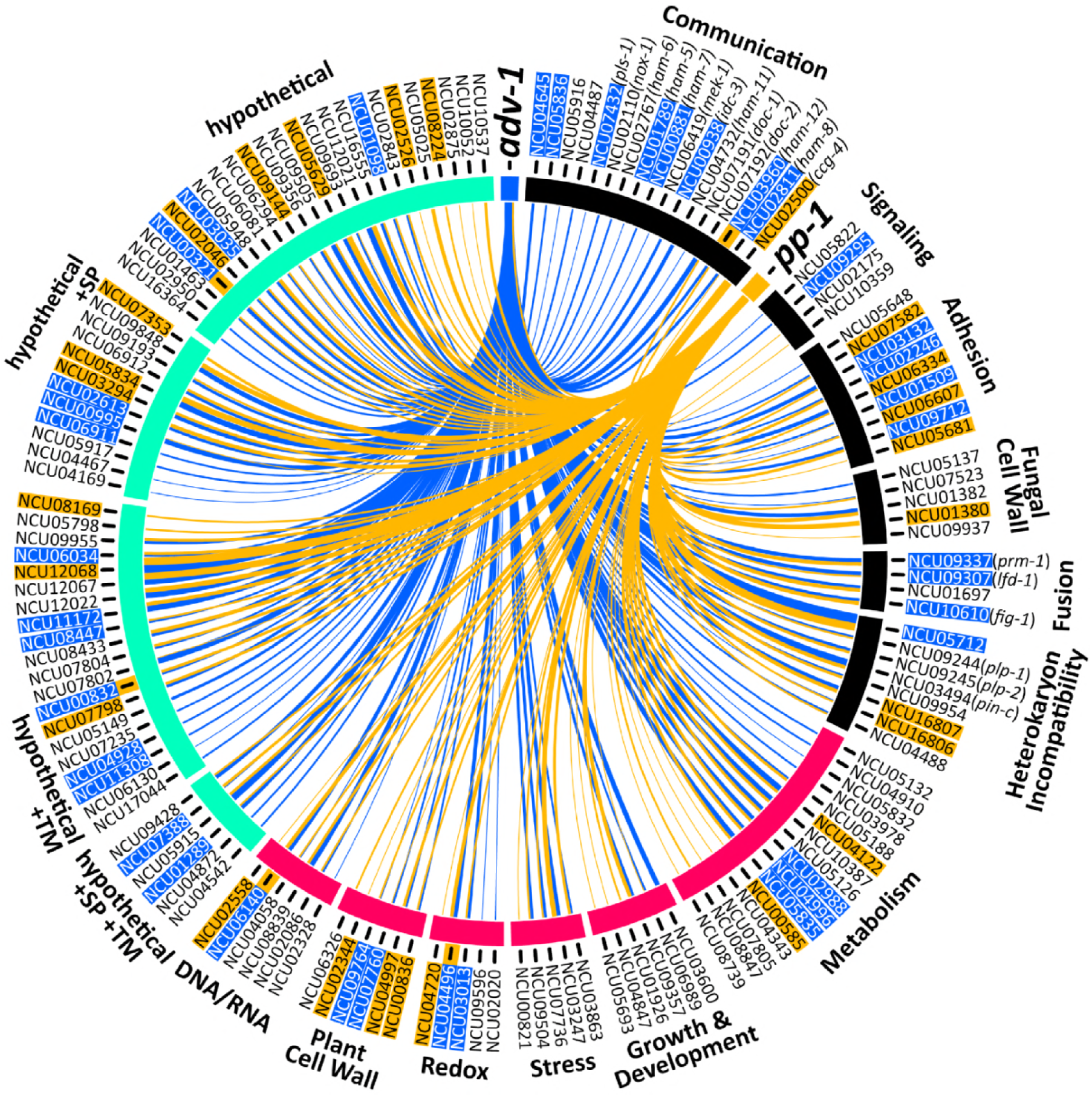
Genes that are positively regulated by ADV-1 and PP-1 in germlings. Circos plots depicting the 155 genes that are significantly down regulated in either *Δadv-1* or *Δpp-1* germlings as compared to parental wild-type germlings (log_2_FC<-2 and p<0.01, consensus among Cuffdiff, DESeq2, and EdgeR). Genes are organized based on function that is either known from previous work, or inferred via homology and protein prediction. Gene function is clustered into three major groups: communication and fusion (black), basic cellular processes (magenta), and hypothetical proteins (seafoam green). Blue lines indicate genes that are regulated by ADV-1, orange lines indicate genes that are regulated by PP-1, and line thickness is proportional to the fold-change difference in expression between the transcription factor mutant and wild type germlings. Gene IDs are highlighted to indicate the presence of at least one ADV-1 (blue) or PP-1 (orange) binding site in the promoter region within 2kb upstream of ATG. Five genes were bound by both ADV-1 and PP-1; these gene IDs are highlighted with blue and have an additional orange highlight immediately adjacent to the gene ID. PP-1 binding sites were determined by consensus between DAPseq and RNAseq, and ADV-1 binding sites are the consensus between DAPseq, RNAseq, and ChIPseq datasets. ChIPseq data is available from Dekhang *et al.* 2017, in which ChIPseq was performed at three different time points. Genes were included here if they were bound by ADV-1 during at least one time point. This list of 155 genes is significantly enriched for ADV-1 binding sites (p=0.0004, Fisher’s Exact Test), but not PP-1 binding sites (p=0.02, Fisher’s Exact Test).

In an independent study, expression patterns in the hyphal stage of the *Δadv-1* mutant were compared to those in wild-type hyphae, at three different circadian time points (Dekhang *et al.* 2017). We compared this *Δadv-1* hyphal dataset with our *Δadv-1* germling dataset (−2<log_2_FoldChange<2, adjusted p-value<0.01, Cuffdiff); all genes differentially expressed during at least one time point were included. We found only a modest overlap (26.5%, 39/147) between our *Δadv-1* germling dataset and the *Δadv-1* hyphal dataset (Figure 1D). Of the 39 genes regulated by ADV-1 in both hyphae and germlings (Table S1), 22 are either predicted or known to be involved in the processes of communication, cell fusion, development, or metabolism including *ham-6, ham-8, ham-11, doc-2, lfd-1, prm-1, mat A-1,* and *esd-C* (Glass *et al.* 1988; Han *et al.* 2008; Fleissner *et al.* 2008; Fu *et al.* 2011; Leeder *et al.* 2013; Palma-Guerrero *et al.* 2014; Heller *et al.* 2016). The remaining 19 genes encode hypothetical proteins. Although the overlap between *Δadv-1* RNAseq datasets can be explained by the fact that chemotropic growth and fusion occur during both germling and hyphal stages, there are additional developmental and morphological differences between germlings and hyphae that could explain reduced overlap between the germling and hyphal *Δadv-1* datasets. Consistent with this hypothesis was the observation that our *Δpp-1* germling dataset overlapped with data from a previous single RNAseq experiment on *Δpp-1* germlings (Leeder *et al.* 2013).

### Identification of ADV-1 and PP-1 binding sites by DAP-seq

DNA-Affinity Purification sequencing (DAP-seq) was recently developed as a high-throughput *in vitro* method for identifying transcription factor binding sites using genomic DNA (O’Malley *et al.* 2016). We adapted this method for *N. crassa*. Briefly, we amplified the *adv-1* and *pp-1* ORFs from cDNA and added a N-terminal HALO-tag to each gene (see Materials and Methods). The HALO-ADV-1 and HALO-PP-1 *in vitro*-synthesized proteins were used for immunoprecipitation experiments using sheared *N. crassa* gDNA that contained adaptors for sequencing. From DAP-seq, we identified PP-1 binding sites that were significantly enriched 2kb upstream of the predicted ATG translation start site for 1953 genes, and ADV-1 binding sites that were significantly enriched within 2kb upstream of the ATG site for 2059 genes (p<0.0001, MACS2)(Figure 3A, File S3).

**Figure 3.**
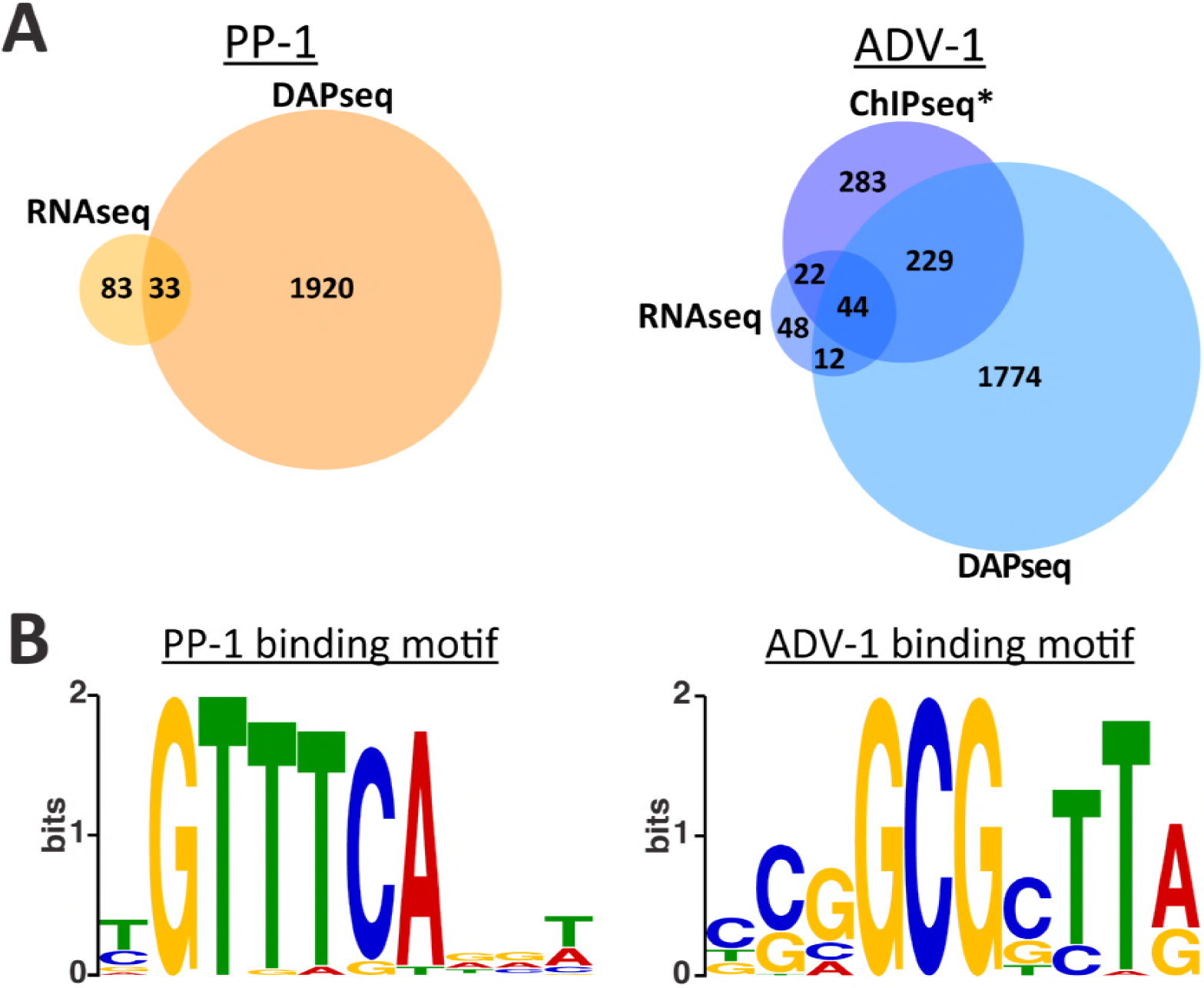
DAPseq identifies promoters bound by ADV-1 or PP-1. **(A)** Number of genes that are down regulated in *Δpp-1* germlings (RNAseq) or bound by PP-1 (DAPseq) (left panel). The number of genes that are down regulated in *Δadv-1* germlings (RNAseq) or bound by ADV-1 (DAPseq and ChIPseq) (right panel). Down regulated genes were identified by consensus between CuffDiff, EdgeR, and DESeq2, log_2_FC<-2 and adjusted p-value<0.01 (compare with Figure 1C). Genes bound by each transcription factor were counted if the transcription factor was bound within 2kb upstream of the ATG (p<0.001). *ADV-1 ChIPseq data is available from Dekhang *et al.* 2017, in which ChIPseq was performed at three different time points. Genes were included here if they were bound by ADV-1 during at least one time point. **(B)** Consensus DNA binding motif for PP-1 or ADV-1 based on DAPseq data.

All *in vitro* assays, such as DAP-seq, contain false positives and false negatives because biologically relevant factors are excluded from the assay. For example, a false negative could arise if a transcription factor requires a co-factor to bind a particular sequence of DNA. A false positive could occur if a factor normally blocks a transcription factor from binding a particular DNA sequence. It is also impossible to know from DAP-seq how environmental or biological conditions affect transcription factor binding. In contrast, *in vivo* methods such as ChIPseq provide a snapshot of where a transcription factor binds under the precise environmental and biological conditions of the assay. Comparing data from both DAP-seq and ChIPseq experiments eliminates some of the error associated with each assay. Additional comparison with expression datasets such as RNAseq can elucidate true-positive peaks associated with genes that are directly bound and regulated by a transcription factor.

Of the 2059 genes bound by ADV-1 identified with DAP-seq, 273 genes were in consensus with a previous ADV-1 ChIPseq dataset (Dekhang *et al.* 2017). Furthermore, 44 of these 273 genes were differentially expressed in *Δadv-1* germlings versus wild type germlings (Figure 2A). The ADV-1 germling regulon (126 genes)(Figure 2) was enriched for ADV-1 binding sites (consensus between ChIPseq and DAP-seq datasets) as compared to ADV-1 binding sites across the entire genome; 35% of the ADV-1 regulon versus 21% of the genome (p=0.0004, Fisher’s Exact Test). Of the 1953 genes bound by PP-1, only 33 genes are differentially expressed in *Δpp-1* vs. wild-type germlings. These 33 genes bound by PP-1 represented 28% of the PP-1 germling regulon (Figure 2), which is what we would expect by chance as compared to the 20% of the entire genome bound by PP-1 (p=0.02, Fisher’s Exact Test).

To construct biologically meaningful transcription factor DNA binding motifs, we analyzed only the strongest DAP-seq peaks that were within a 2kb region upstream of differentially expressed genes. These true-positive peaks were enriched for consensus binding motifs (Figure 3B). The ADV-1 binding motif mirrored previously reported ADV-1 binding motifs (Weirauch *et al.* 2014; Dekhang *et al.* 2017). In contrast, our PP-1 motif differed from the PP-1 motif identified using peptide binding arrays (Weirauch *et al.* 2014).

### PP-1 and ADV-1 regulate genes required for communication, fusion, growth, and development

The *Δadv-1* and *Δpp-1* mutants are impaired in several morphological processes including conidial germination, germling communication, growth rate, aerial hyphae extension, female sexual development, and ascospore germination (Fu *et al.* 2011; Leeder *et al.* 2013; Dekhang *et al.* 2017) (Figure 4, Figure S3, and Table S2). The pleiotropic nature of *Δpp-1* and *Δadv-1* mutants was reflected in the broad functional diversity of the 155 genes that were positively regulated by ADV-1 and/or PP-1 (Figure 2, File S2). These 155 genes fell into three major groups: basic cellular processes (44 genes), communication/fusion/non-self recognition (49 genes), and hypothetical proteins (62 genes). ADV-1 and PP-1 regulate at least some genes in each group, and no group was uniquely regulated by one transcription factor (Figure 2). The significant reduction in growth rate and aerial hyphae in *Δadv-1* and *Δpp-1* cells (Figure 4B,C) can be at least partially explained by the 44 genes whose function was implicated in basic cellular processes such as metabolism, nutrition, growth, development, and general stress response (Figure 2, File S2). Of particular interest to this study are the 49 genes involved in the process of communication, adhesion, fusion, and non-self recognition (Figure 2, File S2). Twenty-two of these 49 genes have been reported previously as being required for normal germling communication, cell fusion, or non-self recognition via heterokaryon incompatibility (Kaneko *et al.* 2006; Kim and Borkovich 2006; Fleissner *et al.* 2008; Fu *et al.* 2011; Leeder *et al.* 2013; Palma-Guerrero *et al.* 2014; Hernández-Galván *et al.* 2015; Lalucque *et al.* 2016; Heller *et al.* 2016, 2018). The remaining 27 genes either have predicted protein domains or homology with proteins implicated in the processes of signaling, adhesion, fusion, cell wall remodeling, or heterokaryon incompatibility.

**Figure 4.**
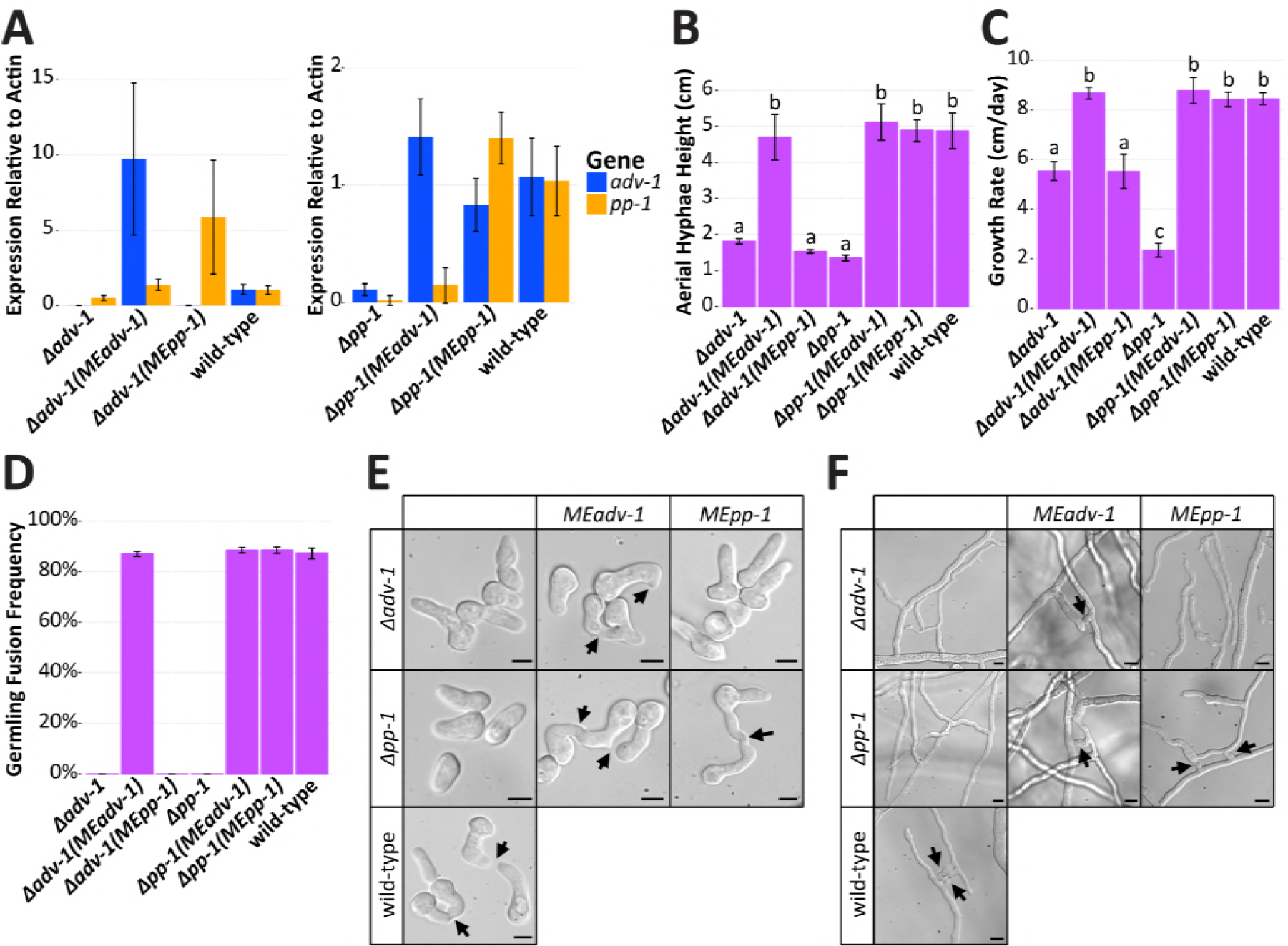
Misexpression of *adv-1* suppresses the phenotype of the *Δpp-1* mutant. **(A)**qRT-PCR data showing mRNA expression levels of *adv-1* and *pp-1* in *Δadv-1 (Ptef1-adv-1-v5; his-3* (*MEadv-1*)) and *Δadv-1 (Ptef1-pp-1-v5; his-3* (*MEpp-1*)) germlings (left panel) and in *Δpp-1* (*Ptef1-adv-1-v5; his-3* (*MEadv-1*)) and *Δpp-1 (Ptef1-pp1-v5;his-3* (*MEpp-1*)) germlings (right panel) compared to the wild type parental strain. **(B)** Mean height of aerial hyphae of strains in (A) three days after inoculation (ANOVA+TukeyHSD, p<0.0001, n=6). **(C)** Mean growth rate per 24 hrs of strains in (A) measured over 4 days (ANOVA+TukeyHSD, p<0.01, n=3). **(D)** Mean frequency of communication and fusion between pairs of germlings for each strain in (A) (n=3, 400-700 germling pairs counted per sample). For all bar plots, error bars indicate standard deviation. **(E)** Photos of the germlings (scale bars = 5µm) and **(F)** hyphae (scale bars = 10µm) for each strain in (A). Arrows indicate chemotropic interactions and successful cell fusion.

We assessed the germling fusion phenotype for each differentially expressed gene that had an available deletion mutant and that had not previously been described (110 deletion mutants available, 23 mutants not unavailable or heterokaryotic, and 22 mutants previously described). Qualitatively, 106 of these mutants had a wild-type-like germling fusion phenotype. However, four mutants (ΔNCU04645, ΔNCU05836, ΔNCU05916, and ΔNCU04487) had obviously reduced levels of germling fusion. We therefore confirmed that the germling fusion phenotype was due to the deleted gene by co-segregation analysis (data not shown), and we quantified the frequency of germling fusion in each strain (Figure 5A). All four mutants have a significantly reduced frequency of germling fusion compared to the wild-type parental strain (p<0.01, ANOVA+TukeyHSD, n=3, ∼200-400 germling pairs per sample). ΔNCU04645 was the only mutant that completely lacked any germling communication or fusion. Similarly, ΔNCU04645 was also the only mutant with significantly reduced height of aerial hyphae (Figure 5B, ANOVA+TukeyHSD, p<0.0001, n=6)(Figure 5B, S4) and growth rate compared to the wild-type parental strain (Figure 5C, ANOVA+TukeyHSD, p<0.01, n=2)(Figure 5C). NCU05836 is predicted to be an ER mannosidase and NCU05916 is a predicted alpha-1,3-mannosyltransferase with homology to the Cu-responsive virulence protein CMT1 in *Cryptococcus neoformans* (Ding *et al.* 2013). Both NCU04487 and NCU04645 are uncharacterized hypothetical proteins. The NCU04487 protein sequence has one predicted C-terminal transmembrane domain and no characterized orthologs. The NCU04645 protein sequence contains a predicted C-terminal AIM24 domain (E-value=1.9E-51, pfam). The AIM24 domain is named after the *S. cerevisiae* protein Aim24p, which is a non-essential mitochondrial inner-membrane protein. In *S. cerevisiae, Δaim24* mutant shows decreased growth, and Aim24p coordinates the assembly of the MICOS (mitochondrial contact site and cristae organizing system) protein complex (Harner *et al.* 2014).

**Figure 5.**
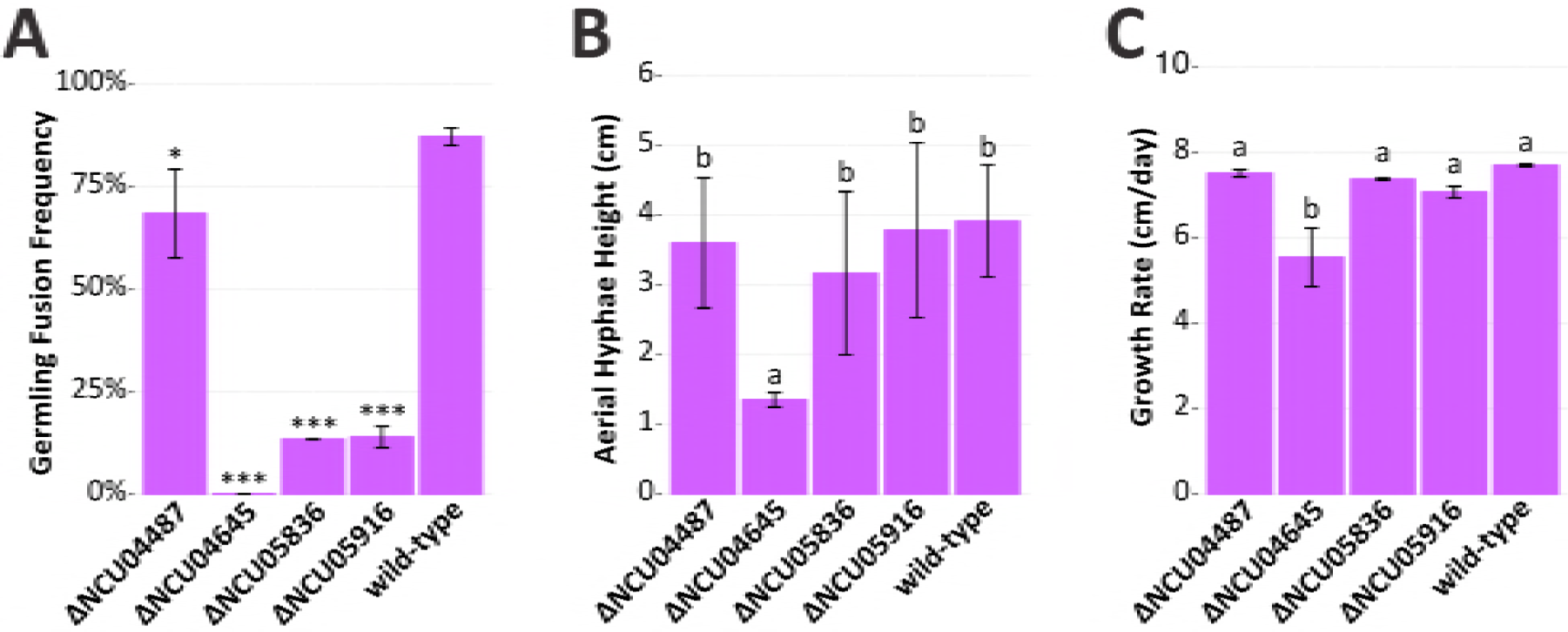
Four ADV-1-regulated genes are required for normal germling fusion and growth. A screen of 110 deletion mutants of the 155 genes regulated by ADV-1 and/or PP-1 revealed that strains carrying mutations in four genes have cell fusion defects. The remaining 151 mutants either have a previously described germling fusion defect, wild-type like germling fusion phenotype, or homokaryotic deletion mutants are not available in the deletion collection. **(A)** ΔNCU04487, ΔNCU04645, ΔNCU05836 and ΔNCU05916 mutants showed reduced germling fusion. Mean germling fusion frequency of each mutant and wild type (n=3, ∼200-400 germling pairs counted per sample). Stars indicate a significant difference compared to the parental wild type strain (*p=0.007,***p<1E-7, ANOVA+TukeyHSD). **(B)** Mean height of aerial hyphae of ΔNCU04487, ΔNCU04645, ΔNCU05836 and ΔNCU05916 mutants three days after inoculation (ANOVA+TukeyHSD, p<0.0001, n=6). **(C)** Mean growth rate of ΔNCU04487, ΔNCU04645, ΔNCU05836 and ΔNCU05916 mutants per day measured over 4 days (ANOVA+TukeyHSD, p<0.01, n=2). For all bar plots, error bars indicate standard deviation.

### Misexpression of *adv-1* suppresses the pleiotropic phenotype of the *Δpp-1* mutant

There are four lines of evidence indicating that PP-1 activates transcription of *adv-1,* with ADV-1 as the primary regulator of the PP-1/ADV-1 regulon. First, *adv-1* is down regulated in the *Δpp-1* mutant, but the inverse is not true. Second, our list of 155 ADV-1/PP-1 regulated genes is significantly enriched for ADV-1 binding sites (p=0.0004, Fisher’s Exact Test), whereas the same is not true for PP-1 binding sites (p=0.02 Fisher’s Exact Test). Third, co-regulated genes are more strongly down regulated in the *Δadv-1* mutant than the *Δpp-1* mutant (Figure 2, File S2). Fourth, there are no potential ADV-1 binding motifs upstream of *pp-1,* but there are several potential PP-1 binding motifs upstream of *adv-1*, despite the fact that DAP-seq did not identify PP-1 bound to the *adv-1* promoter. These observations led to the hypothesis that PP-1 regulates *adv-1* and ADV-1 directly regulates downstream gene expression.

To test this hypothesis, we made epitope-tagged misexpression (ME) constructs *(Ptef1-adv-1-v5* and *Ptef1-pp-1-v5)* and transformed them into *Δpp-1* and *Δadv-1* cells. Successful misexpression in each mutant was verified by qRT-PCR (Figure 4A). As predicted, expression of *adv-1* was restored in *Δpp-1*(*Ptef1-pp-1-v5)* germlings as compared to the *Δpp-1* mutant itself (p=1.8e^−7^, Welch’s t-test, n=8). In contrast, expression levels of *pp-1* were equivalent between *Δadv-1*(*Ptef1-adv-1-v5)* and *Δadv-1* germlings (p=0.0045, Welch’s t-test, n=4) (Figure 4A). Both the *Δadv-1* mutant and the *Δpp-1* mutant have a pleiotropic phenotype when compared with wild-type cells (Figure 4, Figure S3, Table S2). The introduction of *Ptef1-adv-1-v5* fully complemented the pleiotropic phenotype of both *Δadv-1* and *Δpp-1* mutants, while the introduction of *Ptef1-pp-1-v5* only complemented the phenotype of the *Δpp-1* mutant, but not the *Δadv-1* mutant (Figure 4 and Figure S3). Collectively, these misexpression experiments support the RNAseq data and the hypothesis that PP-1 controls expression of *adv-1*; ADV-1 is the primary transcriptional regulator of genes involved in cell communication, fusion, protoperithecial development, and growth.

### PP-1 binds the promoter of *adv-1*

Given that PP-1 was required for expression of *adv-1*, and misexpression of *adv-1* was sufficient to suppress the phenotype of *Δpp-1* cells, we were surprised that our DAP-seq data failed to identify PP-1 binding to the promoter of *adv-1*. We used the consensus DNA binding motif for each transcription factor (Figure 3B) to search for potential binding sites 2kb upstream of both *adv-1* and *pp-1*. Potential ADV-1 binding sites were not identified in the promoter region of either *adv-1* or *pp-1*, however several potential PP-1 binding sites were identified within 2kb of the *adv-1* ORF (Figure 6A). Using antibodies to V5 or GFP, PP-1-bound chromatin was immunoprecipitated from *Δpp-1(Ptef1-pp-1-v5)* and *Δpp-1(Pccg1-pp-1-gfp)* germlings. Ten different primer sets in the *adv-1* promoter region were used to interrogate this pool of PP-1-bound DNA. One primer set successfully amplified a region of DNA ∼500bp upstream of the *adv-1* ORF (Figure 6B & Figure S5). This immunoprecipitated and amplified region is 127bp long and contains two potential PP-1 binding sites (Figure 6C). These data confirm our hypothesis that PP-1 binds the promoter and regulates expression of *adv-1*.

**Figure 6.**
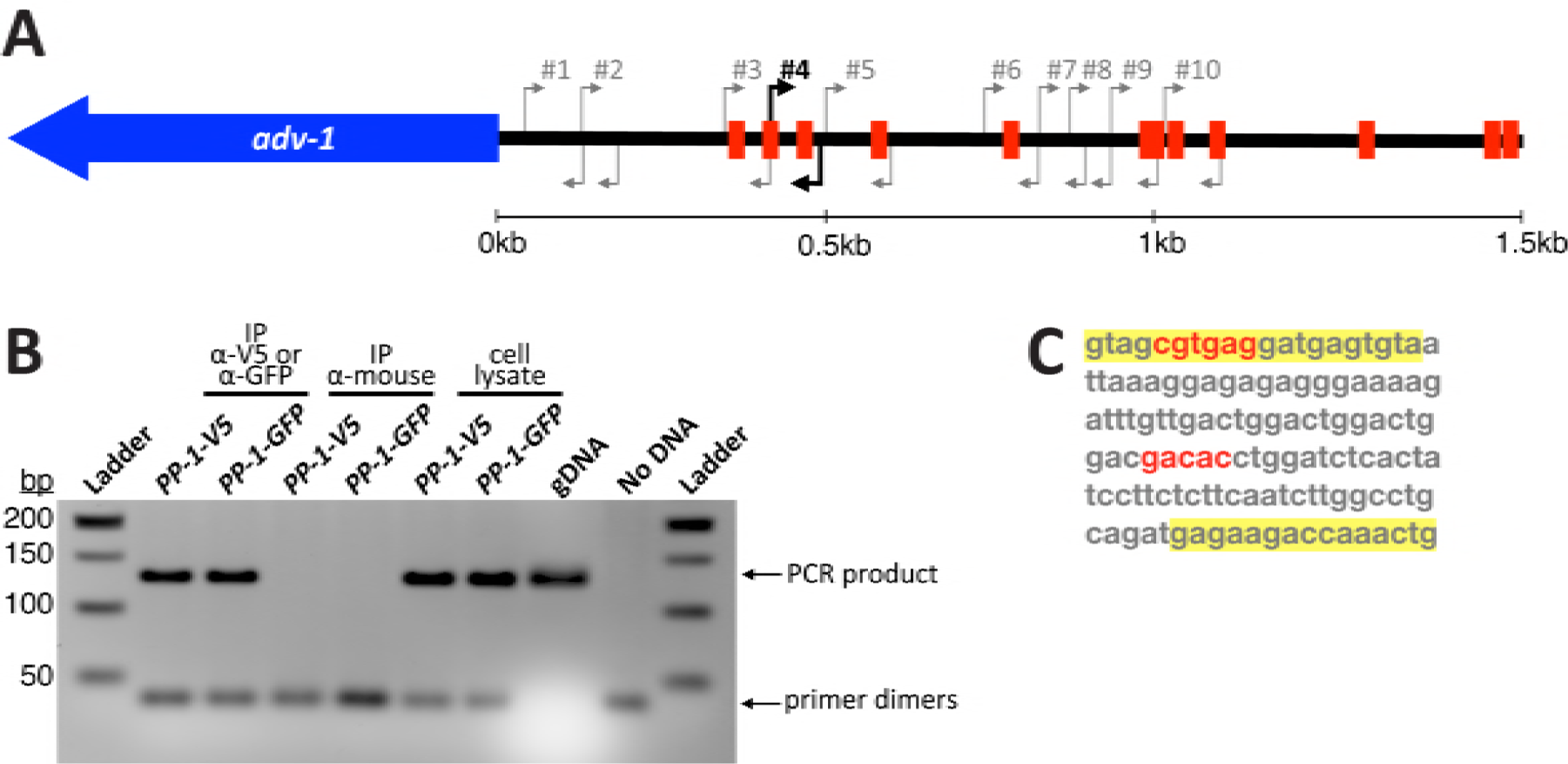
ChIP-PCR identifies a PP-1 binding site ∼500bp upstream of the *adv-1* ORF. **(A)** Diagram of predicted PP-1 binding sites within 1.5kb upstream of *adv-1* ORF, based on the motif depicted in Figure 2B. Arrows indicate ten different PCR primer sets used to interrogate immunoprecipitated chromatin. **(B)** Agarose DNA gel showing results of ChIP-PCR with primer set #4. The remaining PCR results are included in Figure S5. Immunoprecipitation with *Δpp-1(Ptef1-pp-1-v5; his-3)* and *Δpp-1(Pccg1-pp-1-gfp; his-3)* strains was performed using α-V5 or α-GFP antibodies. α-mouse antibodies were used as a negative control. Whole cell lysate and independent *N. crassa* genomic DNA were included as positive PCR controls, with a PCR reaction lacking DNA as an additional negative control. **(C)** The sequence of the PCR product in (B). Primers highlighted with yellow correspond to primer set #4 in (A), predicted PP-1 binding sites are in red.

### Misexpression of *adv-1* rescues growth defects of *Δmak-1*, but not *Δmak-2* mutants

We reasoned that since misexpression (ME) of *adv-1* complemented the phenotype of *Δpp-1*, then *MEadv-1* or *MEpp-1* might also complement the phenotype of MAPK mutants predicted to function upstream of these transcription factors. Previous data indicates that the MAK-1 and MAK-2 MAPK pathways engage in crosstalk and likely function upstream of both PP-1 and ADV-1 (Maerz *et al.* 2008; Maddi *et al.* 2012; Dettmann *et al.* 2012; Leeder *et al.* 2013; Fu *et al.* 2014). In an effort to elucidate this MAPK-transcription factor network, we misexpressed *adv-1* or *pp-1* in the terminal MAPK mutant (*Δmak-1* and *Δmak-2*) for each MAPK pathway; misexpression was confirmed by qRT-PCR (Figure 7A).

**Figure 7.**
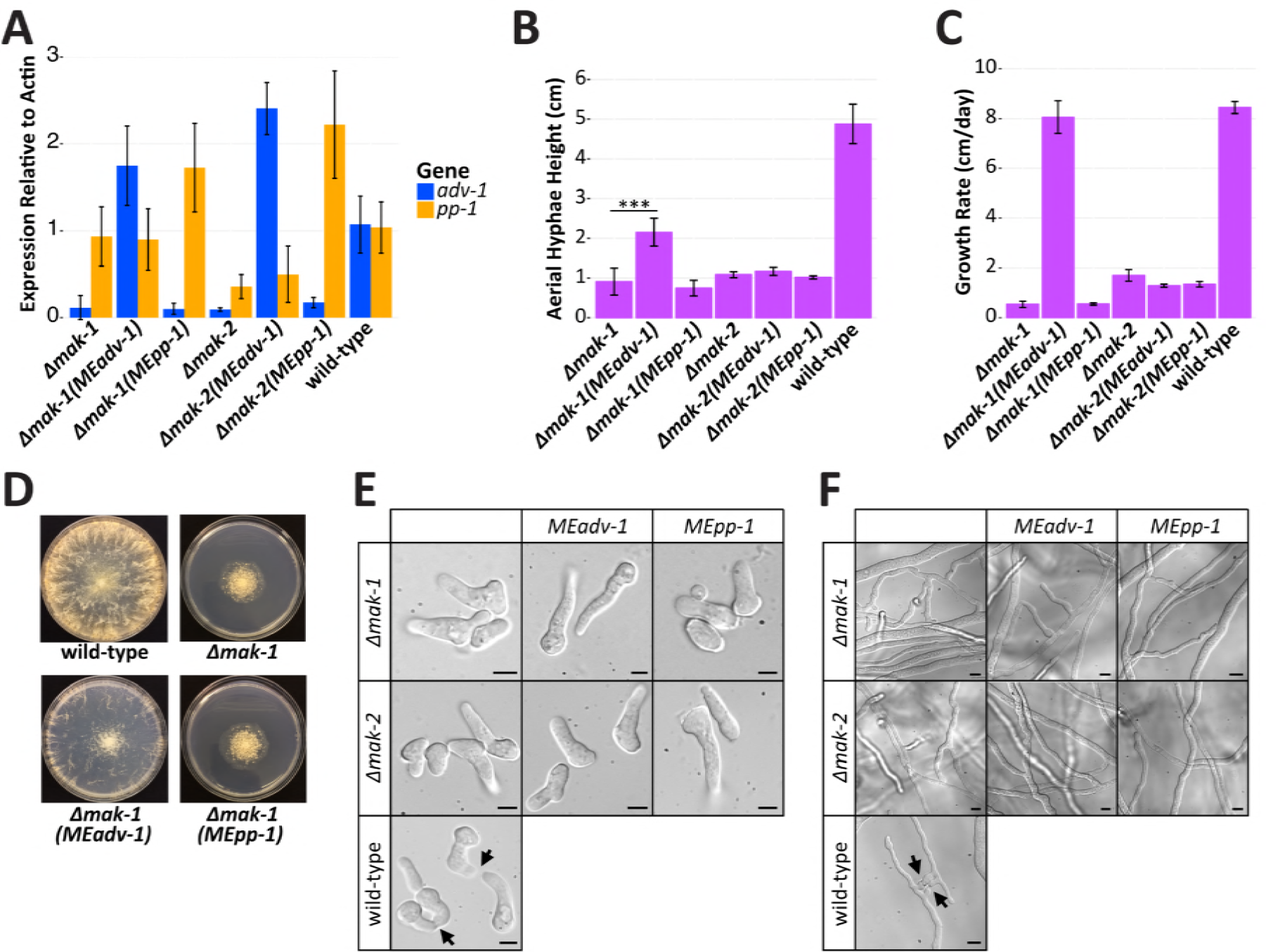
Misexpression of *adv-1* suppresses the growth phenotype of the *Δmak-1* mutant. **(A)** qRT-PCR data showing mRNA expression levels of *adv-1* and *pp-1* in *Δmak-1 (Ptef1-adv-1-v5; his-3* (*MEadv-1*)), *Δmak-1 (Ptef1-pp-1-v5; his-3* (*MEpp-1*)), *Δmak-2 (Ptef1-adv-1-v5; his-3* (*MEadv-1*)) and *Δmak-2 (Pccg1-pp-1-gfp; his-3* (*MEpp-1*)) strains compared to *Δmak-1, Δmak-2* and wild type cells. **(B)** Mean height of aerial hyphae of strains in (A) three days after inoculation (ANOVA+TukeyHSD, ***p=3.6E-8, n=6). **(C)** Mean growth rate per day of strains in (A) measured over 4 days (ANOVA+TukeyHSD, p<0.01, n=3). For all bar plots, error bars indicate standard deviation. **(D)** Colony morphology of the *Δmak-1* mutant relative to the wild-type strain and the *Δmak-1 (Ptef1-adv-1-v5; his-3* (*MEadv-1*)) and *Δmak-1 (Ptef1-pp-1-v5; his-3* (*MEpp-1*)) strains. **(E)** Photos showing a lack of germling fusion (scale bars = 5µm) or **(F)** hyphal fusion (scale bars =10µm) for each strains shown in (A). Arrows indicate chemotropic interactions in the wild-type strain.

Expression of *adv-1* was effectively zero in *Δmak-1* and *Δmak-2* germlings as compared to wild-type germlings (p=3.9e^−19^ and p=1.4e^−23^ respectively, Welch’s t-test, n=8), while expression of *pp-1* was reduced only in *Δmak-2* cells (p=1.15e^−9^, Welch’s t-test, n=8) (Figure 7A). These data indicate that MAK-2 functions upstream of both *pp-1* and *adv-1*, while MAK-1 functions upstream of *adv-1* independently of *pp-1*. To test the effect of misexpressing *pp-1* in *Δmak-2* cells, we used a *Δmak-2(Pccg1-pp-1-gfp)* strain; attempts to obtain *Δmak-2(Ptef1-pp-1-v5)* transformants were unsuccessful. The expression of *Pccg1-pp-1-gfp* complemented the germling communication, cell fusion, aerial hyphae, and growth rate phenotype of *Δpp-1* cells, but not the sexual defects of the *Δpp-1* mutant (Figure S3)(Leeder *et al.* 2013). In contrast to the restoration of *adv-1* expression in *Δpp-1*(*Ptef1-pp-1-v5)* germlings, neither *Δmak-2*(*Pccg1-pp-1gfp)* nor *Δmak-1(Ptef1-pp-1-v5)* germlings showed increased expression of *adv-1* as compared to *Δmak-2 or Δmak-1* germlings, respectively (p=0.73 and 0.06 respectively, Welch’s t-test, n=8). These data indicate that MAK-1 and MAK-2 are necessary for PP-1 dependent transcription of *adv-1*.

We next assessed the phenotypes of each misexpression mutant. In contrast to *Δpp-1*(*Ptef1-adv-1-v5*), *Δpp-1*(*Ptef1-pp-1-v5*), or *Δpp-1*(*Pccg1-pp-1-gfp*) cells, misexpression of *Ptef1-adv-1-v5* or *Pccg1-pp-1-gfp* did not affect on the phenotype of the *Δmak-2* mutant (Figures 7, S3, S6). The misexpression of *Ptef1-pp-1-v5* also had no affect on the phenotype of the *Δmak-1* mutant. However, the misexpression of *Ptef1-adv-1-v5* significantly affected the growth phenotype of the *Δmak-1* mutant (Figure 7B-D). The introduction of *Ptef1-adv-1-v5* into *Δmak-1* cells was sufficient to fully rescue the growth rate defect of the *Δmak-1* mutant (Figure 7C). The *Δmak-1(Ptef1-adv-1-v5*) strain also produced significantly taller aerial hyphae than the *Δmak-1* mutant itself (p=3.6×10^−8^, Welch’s t-test, n=6) (Figure 7B), and misexpression of *Ptef1-adv-1-v5* was sufficient to rescue the compact, rosette-like colony morphology of the *Δmak-1* mutant (Figure 7D). However, the introduction of *Ptef1-adv-1-v5* was insufficient to rescue the communication and cell fusion defects of the *Δmak-1* mutant (Figure 7E, F), including protoperithecial formation and perithecial development (Figure S6). Together these data indicate that growth defects of the *Δmak-1* mutant can be explained by a simple lack of ADV-1. However, for cell-to-cell communication, cell fusion, and sexual reproduction, MAK-1 is clearly necessary, even in the presence of misexpressed *adv-1*.

### *Δpp-1, Δmak-2,* and *Δmak-1* cells are sensitive to cell wall stress, and misexpression of *adv-1* or *pp-1* increases resistance in sensitive strains

MAK-1 is the terminal MAP kinase in the Cell Wall Integrity (CWI) pathway, which maintains the cell wall during growth and in response to stress (Park *et al.* 2008; Maddi *et al.* 2012). Previous studies demonstrated that the *Δmak-1* mutant and other mutants in the CWI pathway are sensitive to the cell wall targeting drugs caspofungin and calcofluor white (Maddi *et al.* 2012). We reasoned that if *Ptef1-adv-1-v5* was sufficient to rescue the growth defects of the *Δmak-1* mutant, then misexpression of *Ptef1-adv-1-v5* might also be sufficient to suppress sensitivity to cell wall stress reagents in *Δmak-1* cells. Additionally, since our data suggested that the MAK-1 and MAK-2 pathways function upstream of both ADV-1 and PP-1, we hypothesized that *Δadv-1, Δpp-1*, and *Δmak-2* cells may also be more sensitive to cell wall targeting drugs.

To test these hypotheses, we assessed growth on agar media containing one of three different cell wall stress drugs; the β-1,3-glucan synthase inhibitor caspofungin (CA), and two different anionic dyes that bind chitin and block chitin-glucan cross-linking; calcofluor white (CFW) and congo red (CR). Wild-type and *Δadv-1* cells were mildly sensitive to all three drugs, while *Δpp-1, Δmak-1*, and *Δmak-2* mutants were almost completely unable to grow on all three drugs (Figure 8). Similar to our other growth phenotype data, both the *Δpp-1(Ptef1-pp-1-v5)* and *Δpp-1(Ptef1-adv-1-v5)* strains showed wild-type-like resistance to all three drugs. Additionally, the misexpression of *Ptef1-adv-1-v5* in *Δmak-1* cells increased its resistance to all three drugs, although this effect was more pronounced on CA than on CFW or CR. The misexpression of *Ptef1-pp-1-v5* in *Δmak-1* cells modestly increased resistance on CFW only. Similarly, the misexpression of *Pccg1-pp-1-gfp* in *Δmak-2* cells modestly increased its resistance to all three drugs, while misexpression of *Ptef1-adv-1-v5* in *Δmak-2* cells did not affect resistance. As expected, the introduction of *Ptef1-pp-1-v5* and *Ptef1-pp-1-gfp* into the *Δpp-1* mutant was sufficient to complement the growth defects of the *Δpp-1* mutant on all three drugs (Figure S7). These data indicate that *adv-1* and *pp-1* function downstream of both MAK-2 and MAK-1 to regulate cell-wall stress responsive gene expression.

**Figure 8.**
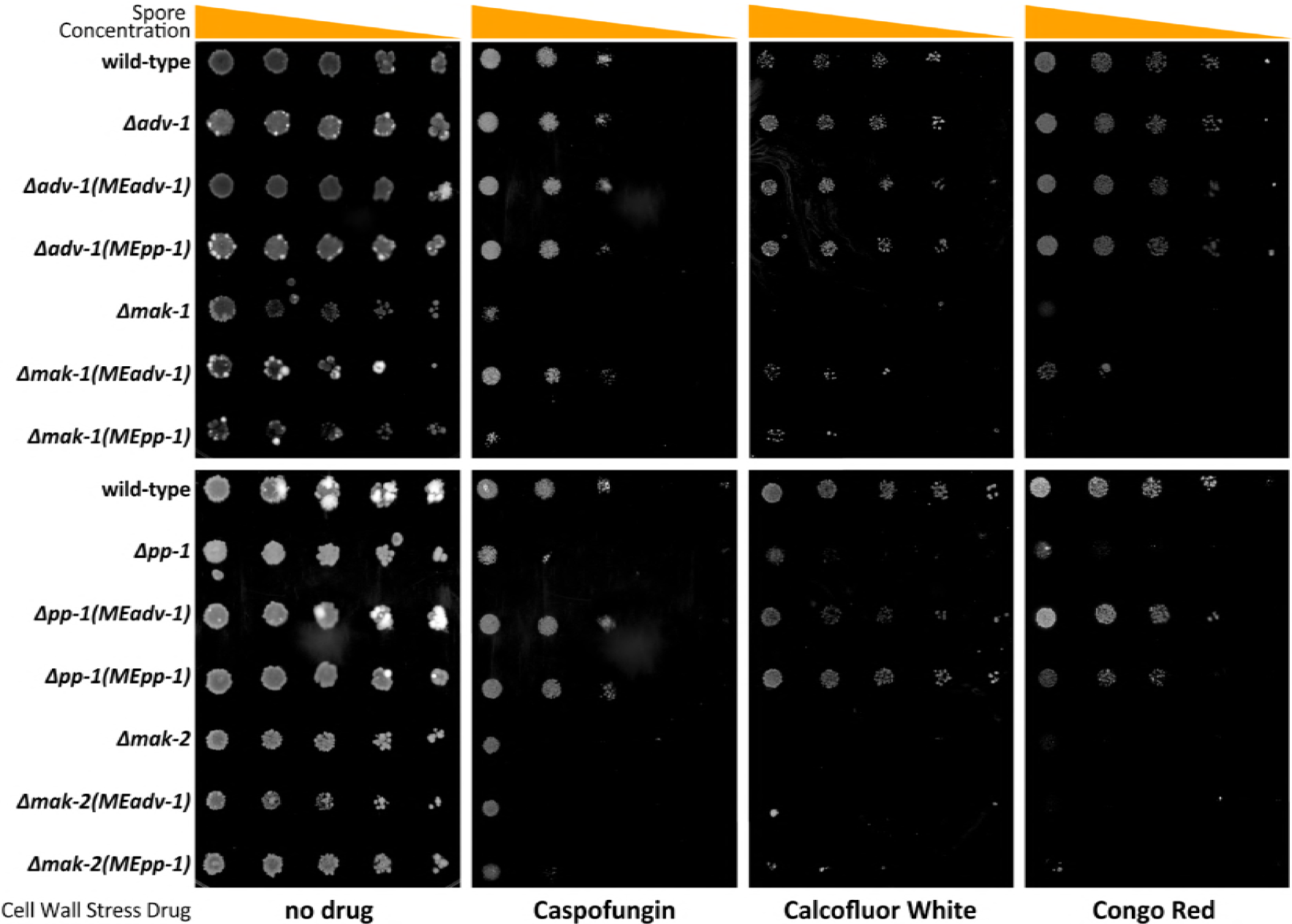
Misexpression of *adv-1* restores resistance to cell wall stress agents in *Δpp-1* and *Δmak-1* cells. A 1:5 serial dilution from ∼5000spores/spot to ∼8spores/spot was performed on *Δpp-1* (*Ptef1-adv-1-v5; his-3* (*MEadv-1*)), *Δpp-1 (Ptef1-pp-1-v5; his-3* (*MEpp-1*)), *Δmak-2 (Ptef1-adv-1-v5; his-3* (*MEadv-1*)), *Δmak-2 (Pccg1-pp-1-gfp; his-3* (*MEpp-1*)), *Δadv-1 (Ptef1-adv-1-v5; his-3* (*MEadv-1*)), *Δadv-1 (Ptef1-pp-1-v5; his-3* (*MEpp-1*)), *Δmak-1 (Ptef1-adv-1-v5; his-3* (*MEadv-1*)) and *Δmak-1 (Ptef1-pp-1-v5; his-3* (*MEpp-1*)) cells compared to *Δpp-1, Δadv-1, Δmak-1, Δmak-2,* and wild-type cells. All agar media contains VMM and FGS to force colonial growth. Plates were incubated at 30°C for 5 days. Drug concentrations: 1.3ug/mL caspofungin,1.5mg/mL calcofluor white, and 1mg/mL congo red.

## DISCUSSION

Our data supports a model for integrated phosphorylation and transcriptional regulation of genes involved in communication, fusion, growth, development, and cell wall stress response by a network of two MAPK pathways and two transcription factors (Figure 9). The MAK-2 pathway primarily functions upstream of PP-1, while the MAK-1/CWI pathway functions upstream of ADV-1. However, PP-1 also directly regulates transcription of *adv-1,* and there are several points of phosphorylation-mediated cross-talk between the MAK-1 and MAK-2 pathways (Maerz *et al.* 2008; Maddi *et al.* 2012; Dettmann *et al.* 2012, 2013; Leeder *et al.* 2013; Fu *et al.* 2014). Additionally, the catalytic activity of MAK-2 is essential for its function (Leeder *et al.* 2013; Jonkers *et al.* 2014). There are seven MAK-2-dependent phosphorylation sites on PP-1 (Jonkers *et al.* 2014), and the catalytic activity of MAK-2 is required for expression of many PP-1 (and ADV-1) regulated genes (Leeder *et al.* 2013). Our qRT-PCR data demonstrated that both MAK-1 and MAK-2 were required for PP-1 dependent transcription of *adv-1* (Figure 7A). Our data also indicate that MAK-1 influences *adv-1*-dependent transcription both directly and indirectly via MAK-2 and PP-1. These data combined with the observation that phosphorylation of MAK-2 is reduced in the *Δmak-1* mutant (Maerz *et al.* 2008; Dettmann *et al.* 2012) indicate that MAK-1 is required for full activation of MAK-2, and that MAK-2 is essential for activating or de-repressing PP-1, likely via phosphorylation. Thus, in the absence of fully-activated MAK-2, misexpression of *pp-1* alone is not sufficient to trigger transcription of *adv-1*. Additionally, our observations that MAK-2 is essential for optimal growth rate and resistance to cell wall stress despite misexpression of *pp-1* or *adv-1* (Figures 7,8) suggests that the catalytic activity of MAK-2 may also be important for post-transcriptional regulation of genes important for growth and cell wall stress response.

**Figure 9.**
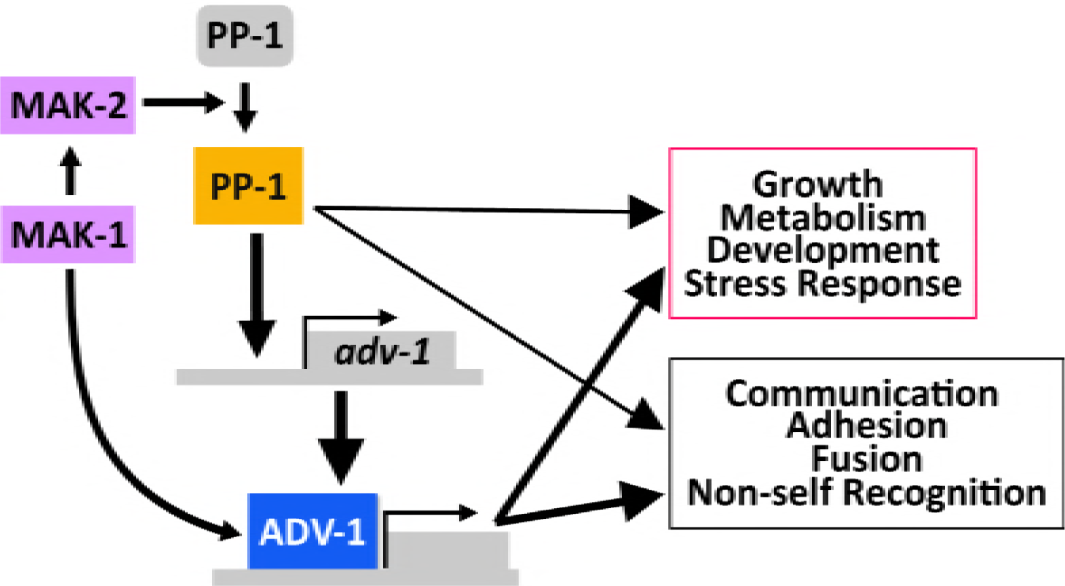
Model for transcriptional regulation by MAK-1, MAK-2, PP-1 and ADV-1. MAK-1 activates both MAK-2 and ADV-1-dependent transcription. MAK-2 activates or de-represses PP-1, which is necessary for transcription of *adv-1* and other down-stream genes (inactivated PP-1 is grey, activated PP-1 is orange). PP-1 directly binds and regulates transcription of *adv-1*; ADV-1 is the direct transcriptional activator of many of the genes required for cell-to-cell communication, cell fusion, growth, development, and metabolism. Additionally, PP-1 and ADV-1 are important for mediating the cell wall stress response downstream of both MAK-1 and MAK-2. While our data indicates that ADV-1 is the primary regulatory for many downstream genes, PP-1 also contributes to the transcription of some of these genes independently of *adv-1*. Downstream gene groups are boxed with colors (magenta or black) that match the colors detailing the same groups in Figure 2.

The proteins that are dependent on MAK-2 for phosphorylation (either directly or indirectly) are enriched for proteins involved in growth, cell cycle progression, development, signaling, and metabolism (Jonkers *et al.* 2014). There are 40 genes that are both phosphorylated in a MAK-2-dependent manner (Jonkers *et al.* 2014) and regulated by ADV-1 or PP-1 (1<log_2_FC<-1, p<0.01, both our data and data from Dekhang *et al.* 2017) (Table S3). These 40 genes include the CWI pathway MAPKK *mek-1,* two hypothetical proteins required for germling communication and fusion (*ham-9* and *ham-11*), one gene that modulates long-distance non-self recognition (*doc-2*), and several genes broadly involved in growth and metabolism. These genes, in addition to the other communication genes that are regulated by ADV-1 and PP-1 may represent a positive feedback loop, in which pathway activation leads to reinforcement. This is particularly true where MEK-1 phosphorylates MAK-1, and then MAK-1 activates transcription of *adv-1*, which in turn activates transcription of MEK-1.

In addition to the communication and fusion genes, there are three calcium signaling genes that are both phosphorylated by MAK-2 (Jonkers *et al.* 2014) and regulated by ADV-1 and PP-1. These genes, a calcineurin binding protein (NCU01504), a calmodulin-dependent kinase (NCU09212), and a predicted Ca^2+^/cation channel (NCU08283) are particularly interesting because calcium is required for polarized growth, cell fusion, and stress response (Silverman-Gavrila 2003; Palma-Guerrero *et al.* 2014, 2015; Virgilio *et al.* 2017). In other systems, the integration of calcium signaling pathways with MAPK pathways has been well documented. For example, in the dicot plant *Nicotiana benthamiana*, ethylene mediates crosstalk between a hormone responsive MAPK pathway and a calcium-signaling pathway (Ludwig *et al.* 2005).

Once activated, the CWI and MAK-2 pathways engage in crosstalk that modulates their responses. The signal(s) and receptor(s) involved in germling communication remain elusive, but it is likely that an unidentified receptor leads to activation of the MAK-2 pathway. An individual receptor may activate both CWI and MAK-2 pathways, similar to how a receptor-associated-Ras-GTPase activates both the ERK pathway (*mak-2* ortholog) and the PI3K-TORC1 MAPK pathway to regulate cell survival, proliferation, and metabolism in mammalian cells (Mendoza *et al.* 2011). Alternatively, there could be more than one receptor such that some receptors lead to activation of the MAK-2 pathway, while other receptors lead to activation of the CWI pathway. The cell wall sensors HAM-7 and WSC-1 both function in the CWI pathway and are required for phosphorylation of MAK-1, and to a lesser extent MAK-2 (Maddi *et al.* 2012). However, *ham-7* is essential for germling fusion while *wsc-1* is dispensable, indicating that these two inputs are differentially interpreted through the CWI pathway. The current model suggests that the MAK-2 pathway is primarily activated or repressed by the sensing of extracellular “self” or “non-self” signals, similar to the orthologous pheromone sensing pathway in yeast (Heller *et al.* 2016). Once activated, the MAK-2 pathway works with the CWI pathway to coordinate growth and prepare for cell fusion by regulating expression of genes involved in adhesion, cell wall remodeling, membrane-merging, and post-fusion non-self recognition. Crosstalk reinforces signaling between these pathways when the integrity of the cell wall is maintained and an extracellular “self” signal is present. However, if cell wall integrity is compromised or if extracellular signaling is dampened, then signaling through both the CWI pathway and the MAK-2 is adjusted to confer the correct response to cell wall stress or termination of communication.

MAP kinases are evolutionarily conserved central regulators for a broad range of cellular processes (Caffrey *et al.* 1999; Xu *et al.* 2016). Crosstalk between MAPK pathways allows for efficient integration of multiple inputs into multiple outputs while also increasing the robustness and plasticity of a signaling network (Jordan *et al.* 2000; Barabási and Oltvai 2004; Komarova *et al.* 2005). In *S. cerevisiae*, the CWI pathway and the osmotic stress (HOG) MAPK pathway respond to environmental stress by integrating input from several different cell-surface sensors via cross-activation and downstream crosstalk, which results in a transcriptional response controlled by several different transcription factors (Rodríguez-Peña *et al.* 2010). Flowering plants, such as *A. thaliana* and *Oryza sativa* have complex MAPK pathways that are characterized by having several modular MAPKKKs and MAPKs for each individual MAPKK (Chardin *et al.* 2017). There are no clear orthologs to the CWI or MAK-2 pathways in plants, however similar signaling networks exist. For example, in *A. thaliana*, crosstalk between MAPK pathways integrates the response to both abiotic and biotic stresses (Fujita *et al.* 2006).

In *N. crassa* and closely related *S. macrospora*, the Striatin-Interacting protein Phosphatase and Kinase (STRIPAK) complex regulates MAPK signaling and mediates some crosstalk between the CWI and MAK-2 pathways. The STRIPAK complex localizes to the nuclear envelope, and mutants in the STRIPAK complex have a similar phenotype to mutants in the CWI or MAK-2 pathways (Simonin *et al.* 2010; Bernhards and Pöggeler 2011; Fu *et al.* 2011; Dettmann *et al.* 2013; Nordzieke *et al.* 2015; Kück *et al.* 2016; Beier *et al.* 2016). Both MAK-1 and MAK-2 have been observed in the nucleus, and the amount of MAK-1 inside the nucleus is dependent on MAK-2 phosphorylating MOB-3, which is a core component of the STRIPAK complex. Furthermore, phosphorylation of MAK-1 is reduced in both the *Δmob-3* mutant and another STRIPAK complex mutant, *Δham-3* (Dettmann *et al.* 2012, 2013). Phosphorylated MOB-3 is also broadly required for fruiting body development, but not chemotropic interactions or STRIPAK complex assembly at the nuclear envelope (Dettmann *et al.* 2013). These data illustrate another potential positive feedback loop in which MAK-1 is required for full phosphorylation of MAK-2, and then MAK-2 phosphorylates MOB-3, which is required for both full phosphorylation of MAK-1 and entry of MAK-1 into the nucleus. Furthermore, the STRIPAK complex clearly mediates CWI-MAK-2 pathway cross talk and modulates the output of both pathways. Future experiments will dissect the interconnected signaling network of the STRIPAK complex, the CWI pathway, and the MAK-2 pathway. Additionally, further characterization of cell-to-cell communication proteins will reveal novel and potentially conserved features of cell-to-cell communication, cell wall dissolution, and membrane merger across the fungal kingdom.

## DATA AVAILABILITY STATEMENT

DAP-seq raw data is available at the NCBI Sequence Read Archive with accession number (SRP133627). RNAseq raw data (.fastq) is available at the NCBI Sequence Read Archive, accession number SRP133239. Differential expression analysis outputs are included with this publication as File S1. File S2 includes detailed information about the 155 differentially expressed genes that we focused on. Results from analyzing DAP-seq data are in File S3.

## ACKNOWLEDGEMENTS

This work was funded by a grant from the National Science Foundation to NLG (MCB1412411). MSF was supported by NIH Genetics training grant #5T32GM007127-40. VWW was supported by NIH Genetics training grant # 5T32GM007127-39. We acknowledge the use of deletion strains generated by Grant P01 GM068087 “Functional Analysis of a Model Filamentous Fungus,” which are publicly available at the FGSC. The RNAseq library prep and sequencing was carried out at the DNA Technologies and Expression Analysis Cores at the UC Davis Genome Center, supported by NIH Shared Instrumentation Grant 1S10OD010786-01.

